# *In Silico* identification of potential drug targets by subtractive genome analysis of *Enterococcus faecium DO*

**DOI:** 10.1101/2020.02.14.948232

**Authors:** Marwah Karim, MD Nazrul Islam, G. M. Nurnabi Azad Jewel

## Abstract

Once believed to be a commensal bacteria, *Enterococcus faecium* has recently emerged as an important nosocomial pathogen worldwide. A recent outbreak of *E. faecium* unrevealed natural and *in vitro* resistance against a myriad of antibiotics namely ampicillin, gentamicin and vancomycin due to over-exposure of the pathogen to these antibiotics. This fact combined with the ongoing threat demands the identification of new therapeutic targets to combat *E. faecium infections*.

In this present study, comparative proteome analysis, subtractive genomic approach, metabolic pathway analysis and additional drug prioritizing parameters were used to propose a potential novel drug targets for *E. faecium strain* DO. Comparative genomic analysis of Kyoto Encyclopedia of Genes and Genomes annotated metabolic pathways identified a total of 207 putative target proteins in *E. faecium DO* that showed no similarity to human proteins. Among them 105 proteins were identified as essential novel proteins that could serve as potential drug targets through further bioinformatic approaches; such as-prediction of subcellular localization, calculation of molecular weight, and web-based investigation of 3D structural characterization. Eventually 19 non-homologous essential proteins of *E. faecium DO* were prioritized and proved to have the eligibility to become novel broad-spectrum antibiotic targets. Among these targets aldehyde-alcohol dehydrogenase was found to be involved in maximum pathways, and therefore, was chosen as novel drug target. Interestingly, aldehyde-alcohol dehydrogenase enzyme contains two domains namely acetaldehyde dehydrogenase and alcohol dehydrogenase, on which a 3D structure homology modeling and *in silico* molecular docking were performed. Finally, eight molecules were confirmed as the most suitable ligands for aldehyde-alcohol dehydrogenase and hence proposed as the potential inhibitors of this target.

In conclusion, being human non-homologous, aldehyde-alcohol dehydrogenase protein can be targeted for potential therapeutic drug development in future. However, laboratory based experimental research should be performed to validate our findings *in vivo*.

## Introduction

*Enterococcus faecium* was believed to be a commensal organism of mammalian digestive tracts that produces bacteriocins and therefore, have been widely used for the production of fermented food products, such as cheese and sausage [1]. Certain strains of *E. faecium* were shown to have beneficial or probiotic [2], effects on animal and human health [3, 4]. However, *E. faecium* has undergone a transition from harmless gut commensals and globally emerged as an important multidrug resistant nosocomial pathogen and is the third most common cause of hospital acquired bacteremia [5, 6]. *E. faecium* is associated with urinary tract infections (UTI), peritonitis, endocarditis, infections in indwelling catheters and septicaemia causing high mortality rate hospitalised patients [7, 8].

Currently, there are over 200 publicly available *E. faecium* draft genomes. Among them TX16 (also referred to as DO) was the first *E. faecium* strain to be isolated and sequenced from an individual with endocarditis and has been used in multiple pathogenesis studies representing the majority of clinical strains globally [9, 10]. The *E. faecium DO* genome consists of one chromosome and three plasmids. The chromosome contains 2,698,137 bp with 2,703 protein-coding ORFs, 62 tRNAs, 6 copies of ribosomal rRNA and 32 other non-coding RNAs. The *DO* genome can be further characterized by numerous hyper-variant loci, a large number of IS elements and transposons. The hyper variable nature of the polysaccharide loci raises the possibility that they are involved in biosynthesis of antigenically diverse surface polysaccharides which could help protect *E. faecium* against host immune responses [11]. The treatment of *E. faecium* mostly relies on conventional antibiotic therapy although there is no such evidence that this improves the course of disease. Furthermore, there has been a progressive increase in antibiotic resistance among *E. faecium* over the last 20 years which is also alarming. With the emergence of vancomycin resistance in 1986 enterococci can now be resistant to all currently approved antimicrobial agents [12-14]. The accumulated results have strongly provoked a necessity for developing a potential drugs and vaccine candidates to tackle this dreadful pathogen.

Identification of therapeutic drug targets is the first step in the drug discovery process. Due to the recent advances in complete genome sequencing, bioinformatics, cheminformatics and availability of both pathogen and host-genome sequences, computational methods have been used widely for the identification of potential drug and vaccine targets at genomic levels in different pathogenic microorganisms. Among various strategies available, subtractive and comparative genomics approach combined with metabolic pathway analysis was considered worthy to identify the proteins essential for the pathogen but absent in the host [15-17]. To search for non-human homologous targets, subtraction of the host genome from essential genes of pathogens ensures no interaction of drugs with human targets. On contrary, comparative genomics method emphasizes on selected conserved proteins across different species as most favourable targets [15, 18]. The use of advanced bioinformatics tools with integrated genomics, proteomics, and metabolomics may ensure the discovery of potential drug targets for most of the infectious diseases. Once the target(s) have been identified, it is expected that the identified potential drug and vaccine targets will not only expand our understanding on the molecular mechanisms of *E. faecium DO* pathogenesis but also facilitate the production of novel therapeutics.

In this present study, we performed *in silico* analysis to identify the novel therapeutic targets in *E. faecium DO* by combining analysis of metabolomics and genomics data. Instead of analysing the whole genome, the key essential or survival proteins of the pathogen were considered that are non-homologous to the host. A good number of novel targets in *E. faecium DO* were elucidated to design effective drugs against broad-spectrum pathogenic bacteria. Moreover, we provided a modeled 3D structure and *in silico* docking of aldehyde-alcohol dehydrogenase which was selected as the best possible target for designing potential drugs. Finally, our study proposed suitable ligands which can be potential inhibitors of this enzyme. To the best of our knowledge this is the first computational and subtractive genomics analysis of different metabolic pathways for the identification of potential drug and vaccine targets in *E. faecium DO*.

## Materials and methods

### Comparative analysis of host-pathogen pathways and protein retrieval

A systematic hierarchical workflow to identify and characterize the potential drug targets for *E. faecium DO* were summarized in Figure 1 and sequentially mentioned below. Genome-wide metabolic pathway analysis for both human and *E. faecium DO* was performed using Kyoto Encyclopedia of Genes and Genomes (KEGG) database [19]. The identification numbers of all metabolic pathways from both organisms were extracted from the database. A manual comparison was then conducted by placing the name of individual pathway of pathogen against the pathways of the host, *H. sapiens*. According to the KEGG database annotations, pathways that were absent in the human but did appear in the pathogen were considered as unique to *E. faecium DO*, whereas the remaining pathways were listed as common pathways. Proteins with their corresponding amino acid sequences for both unique and common pathways were obtained from UniprotKB [20].

**Figure 1.**
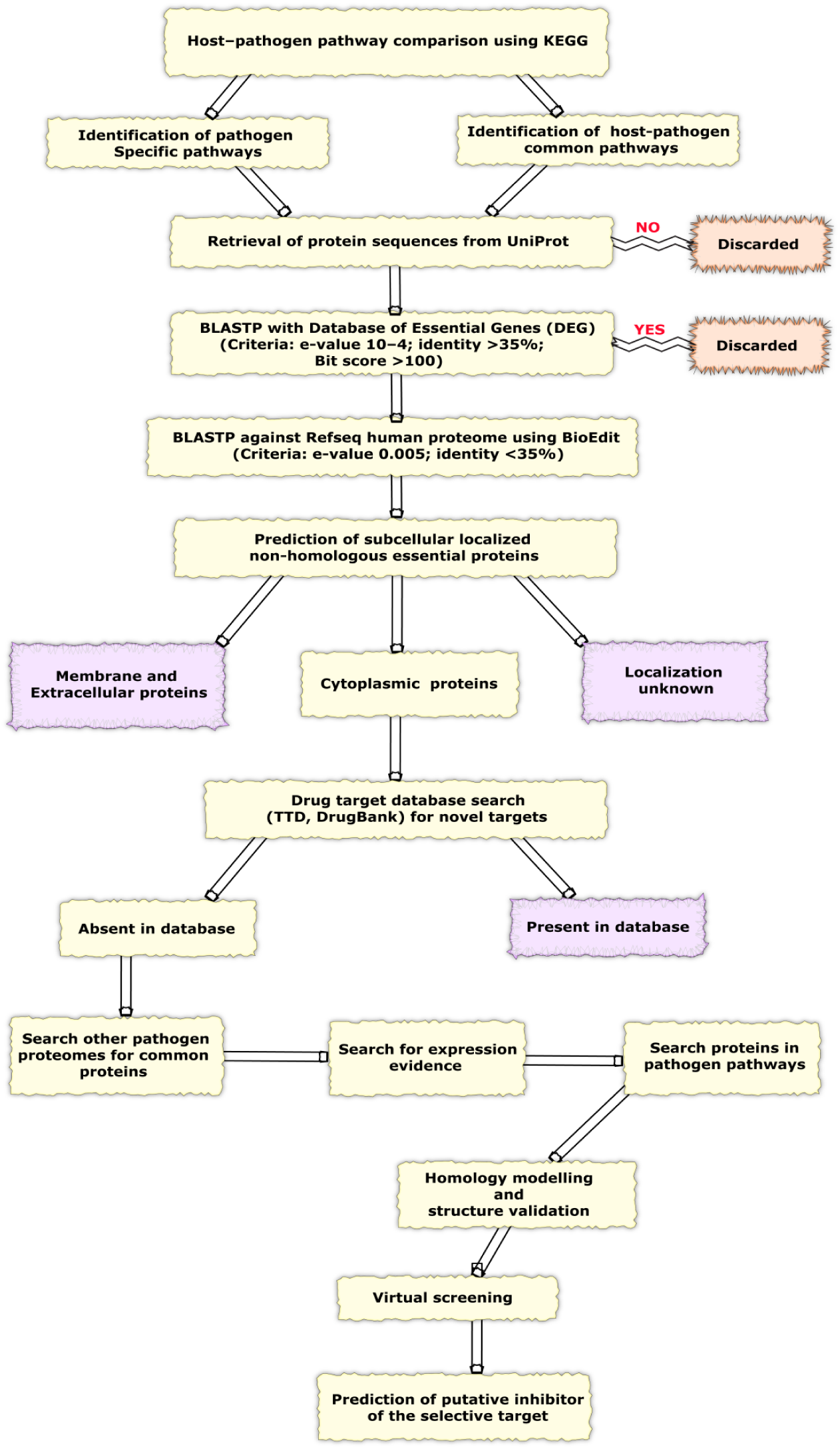
A schematic representation of the workflow of computational drug target identification and prediction of putative inhibitors of the selected target. Abbreviations: BLASTP = Protein Basic Local Alignment Search Tool; TTD = Therapeutic Target Database.

### Identification of essential and non-homologous pathogen proteins

To identify the essential proteins involved in unique and common host-pathogen pathways, the dataset was analyzed by Database of Essential Genes (DEG) version 6.8 [21], with cut off for e-value 10^−4^, sequence identity <35%, bit score >100 and others as default. The obtained essential proteins were subjected to Protein Basic Local Alignment Search Tool (BLASTP) using BioEdit Sequence Alignment Editor (version 7.1.3) against the human protein sequences retrieved from Refseq database. Only the non-hit proteins at e-value cut off 0.005 and <35% identity were selected as non-homologous pathogen proteins to avoid any functional redundancy with host proteome.

### Prediction of subcellular localization and drug targets prioritization

Subcellular localization of the essential non-human proteins was predicted by PSORTb version 3.0.2 [22] and three types of localization such as: cytoplasmic, membrane, and extracellular proteins for Gram-negative bacteria were predicted. Several molecular and structural criteria that have been proposed to aid in prioritizing suitable drug targets [23] were evaluated for each of the predicted drug targets in *E. faecium DO*. This involved, calculation of molecular weight (MW) using computational tools and drug targets associated literature available at Swiss-Prot database [24]. The presence of experimentally and computationally solved 3D structures were detected by searching the Protein Data Bank (PDB) database [25].

### Novel targets identification and hunt for common proteins

To identify novel targets among the potential targets, databases DrugBank, SuperTarget, and Therapeutic Target Database (TTD), were searched for similarity with the cytoplasmic proteins [26, 27]. E-value <10^−5^, sequence identity >35%, and bit score >100 were set as parameters and the non-hit proteins at the threshold value were selected as novel drug targets. In addition, all protein sequences of 73 different strains of pathogenic bacteria were retrieved from PATRIC database with a goal to sort common proteins amongst pathogenic bacteria [28, 29]. The novel targets were subjected to BLASTP against these proteomes at e-value cut off 10^−5^, sequence identity >35%, bit score >100 with BioEdit software. The proteins that were found to be common in at least 20 pathogenic strain proteomes were listed as broad-spectrum targets and different bacterial species were used as references.

### Homology modelling

As no exact PDB structure was available for aldehyde-alcohol dehydrogenase in PDB, therefore it’s two domains-acetaldehyde dehydrogenase and alcohol dehydrogenase were subjected to BLAST search against PDB structures using 0.001 e-value cut off. The template for homology modelling was chosen considering X-ray diffraction resolution and highest sequence similarity using SWISS-MODEL server [30].

### Structure validation and active site prediction

The modelled structure was assessed using SWISSMODEL structure assessment tool [31] and ANOLEA [32] to assess the packing quality of the models. The stereochemical quality of protein structures were checked through PROCHECK suite of programs [33]. Energy minimization was carried out by GROMOS96 with default parameters implemented in Swiss PDB Viewer (version 4.0.4) [34], followed by a CASTp server [35] analysis to determine the active site of the modelled structure.

### Virtual screening, ADME and toxicity analysis

Virtual screening was carried out with a total of 9317 molecules from DrugBank, anti-bacterial drugs of DrugBank [36] and Antimicrobial Drug Database [37] based on selected active sites of both acetaldehyde dehydrogenase and alcohol dehydrogenase domains. Molecules were then converted using Open Babel command line application [37] and Raccoon software [38]. DrugBank molecules were filtered by Raccoon using Lipinski-like options whereas molecules with molecular weight >1000 were discarded for Antimicrobial Drug Database. After filtering, we got 6308 (1436 FDA-approved, 4050 experimental, 415 investigational, illicit178, withdrawn 158 and 71 nutraceutical) and 2882 (1842 antibacterial and 1040 antifungal) molecules from DrugBank and Antimicrobial Drug Database respectively. No filtering was applied for 137 anti-bacterial drugs of DrugBank. Virtual screening was performed using The Texas Advanced Computing Center (TACC) computing resources. Furthermore, Pymol and Discovery Studio (Accelrys, San Diego, CA, USA) were used for protein-ligand interaction analysis and visualization; PoseView was used to generate interaction diagrams. ADMET prediction and Oral bioavailability were done using PreADMET server [39] and FAF-Drugs3 program of Mobyle@RPBS server [40].

## Results

### Identification of pathogen specific and host-pathogen common metabolic pathways

In our study, a list of potential drug and vaccine targets were identified using computational, comparative and subtractive genomics analysis of different metabolic pathways from *E. faecium DO*. A systematic workflow involving several bioinformatics tools, databases, and drug target prioritization parameters (Figure 1) was defined, with an ultimate goal of obtaining information about proteins that were involved in various metabolic pathways of *E. faecium DO*, but absent in its host, therefore, minimizing potential side effects. A total of 108 different metabolic pathways involved in pathogen were obtained from KEGG database. Subsequently, a comparative analysis of the metabolic pathways of the host and pathogen revealed a total of 26 pathogen specific pathways and 82 host-pathogen common pathways (Table 1).

**Table 1.** Host–pathogen common and pathogen specific pathways from KEGG database

| Pathway IDs | Pathway Names | Pathway IDs | Pathway Names |
| --- | --- | --- | --- |
| **Host-Pathogen Common Pathways** |  |  |  |
| efu00010 | Glycolysis /Gluconeogenesis | efu00785 | Lipoic acidmetabolism |
| efu00040 | Pentose andglucuronate interconversions | efu00790 | Folate biosynthesis |
| efu00051 | Fructose andmannose metabolism | efu00900 | Terpenoid backbone biosynthesis |
| efu00052 | Galactose metabolism | efu00910 | Nitrogen metabolism |
| efu00053 | Ascorbate and aldarate metabolism | efu00920 | Sulfur metabolism |
| efu00061 | Fatty acid biosynthesis | efu00970 | Aminoacyl-tRNA biosynthesis |
| efu00130 | Ubiquinone and other terpenoid-quinone biosynthesis | efu01040 | Biosynthesisof unsaturated fatty acids |
| efu00190 | Oxidative phosphorylation | efu01200 | Carbon metabolism |
| efu00220 | Arginine biosynthesis | efu01210 | 2-Oxocarboxylic acid metabolism |
| efu00230 | Purine metabolism | efu01212 | Fatty acid metabolism |
| efu00240 | Pyrimidine metabolism | efu01220 | Degradationof aromatic compounds |
| efu00250 | Alanine aspartate and glutamate metabolism | efu01230 | Biosynthesisof amino acids |
| efu00260 | Glycine serine and threonine metabolism | efu02010 | ABC transporters |
| efu00261 | Monobactam biosynthesis | efu03018 | RNA degradation |
| efu00270 | Cysteine andmethionine metabolism | efu03020 | RNA polymerase |
| efu00280 | Valine leucine and isoleucine degradation | efu03030 | DNA replication |
| efu00290 | Valine leucine and isoleucine biosynthesis | efu03060 | Protein export |
| efu00300 | Lysine biosynthesis | efu03410 | Base excision repair |
| efu00330 | Arginine andproline metabolism | efu03420 | Nucleotide excision repair |
| efu00350 | Tyrosine metabolism | efu03430 | Mismatch repair |
| efu00360 | Phenylalanine metabolism | efu03440 | Homologous recombination |
| efu00400 | Phenylalanine tyrosine and tryptophan biosynthesis | efu04122 | Sulfur relay system |
| efu00410 | beta-Alaninemetabolism | **Pathogen Specific Pathways** |  |
| efu00430 | Taurine andhypotaurine metabolism | efu03070 | Bacterial secretion system |
| efu00450 | Selenocompound metabolism | efu02020 | Two-component system |
| efu00460 | Cyanoamino acid metabolism | efu00903 | Limonene and pinene degradation |
| efu00471 | D-Glutamineand D-glutamate metabolism | efu00642 | Ethylbenzene degradation |
| efu00480 | Glutathionemetabolism | efu00621 | Dioxin degradation |
| efu00500 | Starch and sucrose metabolism | efu00401 | Novobiocin biosynthesis |
| efu00511 | Other glycandegradation | efu00253 | Tetracyclin biosynthesis |
| efu00520 | Amino sugarand nucleotide sugar metabolism | efu01503 | Cationic antimicrobial peptide (CAMP) resistance |
| efu00523 | Polyketide sugar unit biosynthesis | efu01502 | Vancomycin resistance |
| efu00561 | Glycerolipidmetabolism | efu01501 | beta-Lactamresistance |
| efu00562 | Inositol phosphate metabolism | efu01054 | Nonribosomalpeptide structures |
| efu00564 | Glycerophospholipid metabolism | efu00680 | Methane metabolism |
| efu00590 | Arachidonicacid metabolism | efu00643 | Styrene degradation |
| efu00600 | Sphingolipidmetabolism | efu00627 | Aminobenzoate degradation |
| efu00620 | Pyruvate metabolism | efu00626 | Naphthalenedegradation |
| efu00630 | Glyoxylate and dicarboxylate metabolism | efu00625 | Chloroalkaneand chloroalkene degradation |
| efu00640 | Propanoate metabolism | efu00622 | Xylene degradation |
| efu00650 | Butanoate metabolism | efu00550 | Peptidoglycan biosynthesis |
| efu00660 | C5-Brancheddibasic acid metabolism | efu00521 | Streptomycinbiosynthesis |
| efu00670 | One carbon pool by folate | efu00473 | D-Alanine metabolism |
| efu00730 | Thiamine metabolism | efu00440 | Phosphonateand phosphinate metabolism |
| efu00740 | Riboflavin metabolism | efu00362 | Benzoate degradation |
| efu00750 | Vitamin B6 metabolism | efu00361 | Chlorocyclohexane and chlorobenzene degradation |
| efu00760 | Nicotinate and nicotinamide metabolism | efu00332 | Carbapenem biosynthesis |
| efu00770 | Pantothenateand CoA biosynthesis | efu00121 | Secondary bile acid biosynthesis |
| efu00780 | Biotin metabolism |  |  |

### Identification of non-homologous essential proteins

We identified 617 probable essential proteins from host–pathogen common pathways and 143 from pathogen specific unique pathways (Supplementary file 1), which showed good similarity with the experimentally proven essential proteins recorded in DEG database. Screening through BlastP, we found considerable similarities (E-value less than 10^−4^) between pathogen and host proteins which were identified as homologous and excluded. Hits having expectation value greater than 10^−4^ were selected as non-homologous proteins to host. We also found 184 and 23 proteins from host-pathogen common and host-pathogen unique pathways respectively (Supplementary file 2) which are essential for pathogen; however, non homologous to host.

### Subcellular localization and prediction of drug target prioritization

Subcellular localization of proteins in a cell is an important feature that can determine their potential functions, identification of suitable and effective drug targets. Cytoplasmic proteins are more favourable as therapeutic drug targets as the membrane localized proteins are difficult to purify [38]. In addition, identified non-homologous essential proteins of *E. faecium DO* were further characterized based on other essential features, such as; accessibility value of a target protein, preferably low MW (<100 kDa), presence of transmembrane helixes and availability of 3D structural information. It has been demonstrated that proteins with low molecular weight (100–110 kDa) increase the accessibility value of a target protein. In our study, majority of the proteins had MW less than 100 KDa suggesting them as possible candidate for drug development to be studied experimentally.

A total of 184 proteins were found from host-pathogen common pathway whereas 157 proteins were found to be cytoplasmic, 18 proteins to be membrane localized and rest 9 proteins to be of unknown localization. In addition, from 23 proteins from pathogen specific pathways, 17, 4, 2 proteins were found to be cytoplasmic, membrane localized, and of unknown localization respectively (Figure 2 & Supplementary file 3). Based on these results, 157 and 17 cytoplasmic proteins from common pathways and unique pathways respectively were considered for further analysis to identify suitable drug targets.

**Figure 2.**
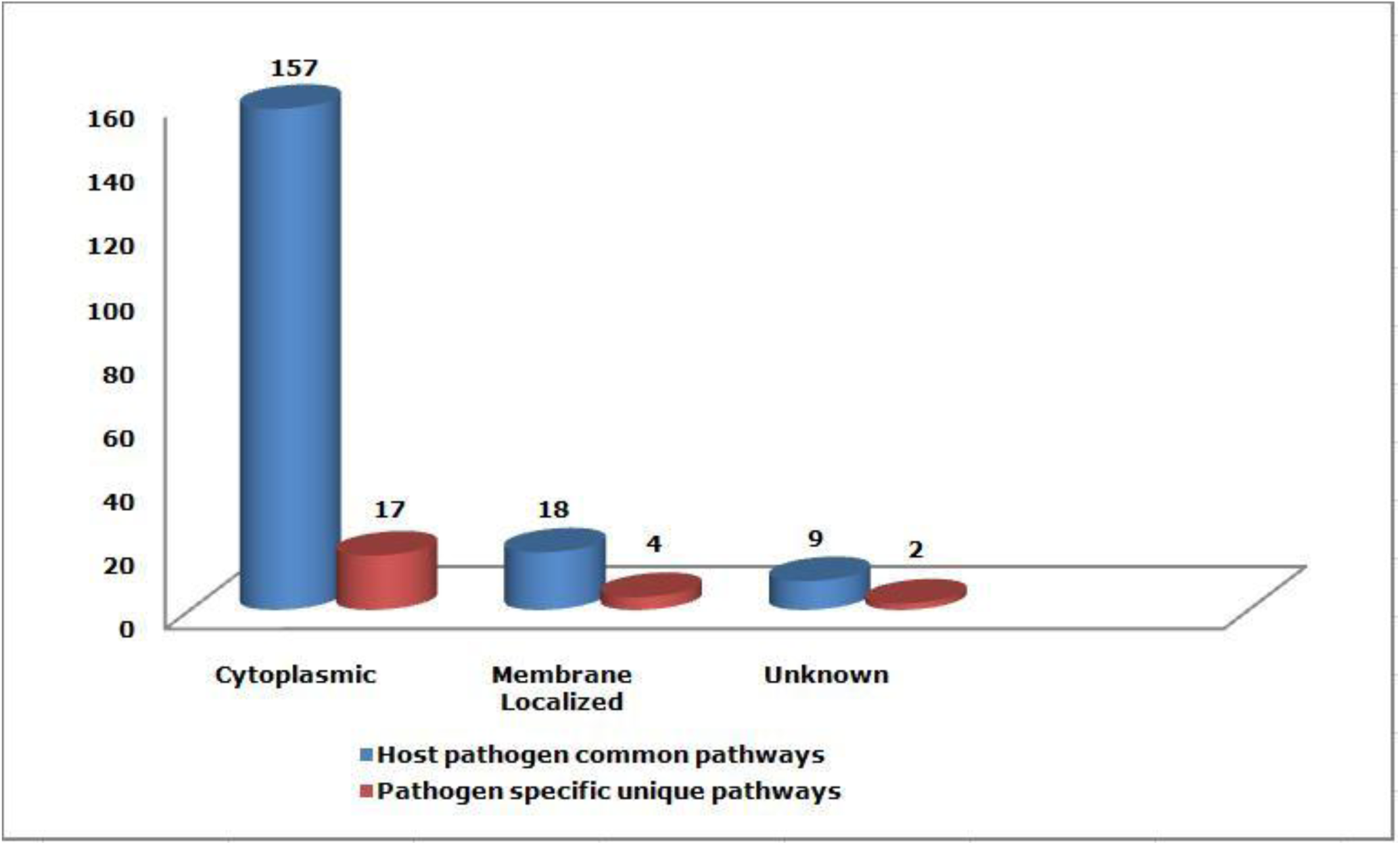
Comparative subcellular localization of proteins from the common host-pathogen pathways and pathogen-specific pathways.

### Novel targets in pathogen unique and host–pathogen common pathways

Proteins which showed significant similarity with the databases were discarded and the remaining protein sequences were taken as novel targets. Our study revealed 9 proteins that were uniquely involved in pathogen specific 6 unique pathways and these are-Bacterial secretion system [PATH: efu03070], Two-component system [PATH: efu02020], Cationic antimicrobial peptide (CAMP) resistance [PATH: efu01503], Methane metabolism [PATH: efu00680], Tetracycline biosynthesis and Naphthalene degradation [PATH: efu00626].

In addition, we identified 96 proteins in 44 host–pathogen common metabolic pathways as novel targets (Supplementary file 4), among them ten protein targets were involved in bacterial secretion system pathway of *E. faecium*. Bacterial system secrete a wide range of proteins whose functions include biogenesis of organelles, bacterial growth and survival in eukaryotic host cells [41]. A total of 17 proteins were identified as unique in the pathogen specific pathway in a two-component system (KEGG Pathway: efu02020). Bacterial two-component systems are signalling pathways that regulate bacterial characteristics such as virulence, pathogenicity, symbiosis, motility, nutrient uptake, secondary metabolite production, metabolic regulation and cell division. It regulates myriad of physiological processes in response to various environmental or cellular parameters enabling bacteria to adapt in adverse conditions. Therefore, they can be considered as potential targets for antimicrobial drug designing [42]. Cationic antimicrobial peptides (CAMPs) are among the most ancient and efficient components of host defence against invasive infections by pathogenic bacteria. CAMPS are produced by epithelial cells and immune cells such as neutrophils and macrophages [43, 44]. Therefore, CAMPs have been considered as promising candidates to treat infections caused by pathogenic bacteria both in animals and humans. *Streptococcus pneumoniae, Streptococcus pyogenes*, and staphylococci, organisms that cause respiratory and cutaneous infections, and members of the *Enterobacteriaceae* and *Pseudomonas families*, organisms that cause diarrhea, urinary infection, and sepsis, are now resistant to virtually all of the older antibiotics such as-beta-Lactam, vancomycin, Streptomycin, Carbapenem etc. AckA was found to be uniquely present in methane metabolism (KEGG Pathway: map00680) pathway. Methanotrophs consume methane as their sole source of carbon and energy for growing whereas methanogens obtain energy for growth by oxidizing a limited number of substrates to methane under anaerobic conditions [45]. Another pathway with significant impact on bacteria follows antibiotic biosynthesis. The clinical importance of *E. faecium* is related to its antibiotic resistance that contributes to the risk of colonization and infection. The presence of tetracyclin biosynthesis pathway in bacteria can yield a large number of biosynthetically and chemically modified variants, most of which have found to be valuable pharmaceutical compounds [46]. Moreover, there are bacteria’s such as Pseudomonas spp., Vibrio spp., Mycobacterium spp., Marinobacter spp., and Sphingomonas spp. which possess naphthalene-degrading pathway. Naphthalene has long been used as a model compound in polycyclic aromatic hydrocarbon (PAHs) bioremediation studies. PAH bioremediation is therefore considered an effective and environmental friendly cleanup strategy as it involves the partial or complete bioconversion of pollutants to microbial biomass, carbon dioxide and water [47].

### Identification of broad-spectrum targets

Proteins that are common across bacterial genera, would be a promising candidate for broad-spectrum antibiotic targets. In this study, 73 bacterial species (Supplementary file 5) were included as reference and proteins which were common in at least 20 different species were listed and 19 proteins were identified as broad-spectrum target proteins (Supplementary file 6). These proteins are involved in multiple regulatory pathways, therefore, would be better targets as inhibition of their activity will impede one or more pathways in the pathogen [48]. Aldehyde-alcohol dehydrogenase was involved in maximum number of pathways. This enzyme has three activities; ADH, ACDH, and PFL-deactivase. In aerobic conditions it acts as a hydrogen peroxide scavenger. The PFL deactivase activity catalyzes the quenching of the pyruvate-formate-lyase catalyst in an iron, NAD, and CoA dependent reaction. There is no resolved X-ray crystallography structure for aldehyde-alcohol dehydrogenase (*E. faecium* DO) as we intended to do homology modeling. Therefore, aldehyde-alcohol dehydrogenase subunit (Uniprot ID: I3TYJ0) was chosen for homology modeling and docking studies. We conducted BLAST searching for aldehyde-alcohol dehydrogenase (Uniprot ID: I3TYJ0) with e-value cut off of 0.001 against UniprotKB. We selected the organisms whose proteins showed at least 77% sequence identity with aldehyde-alcohol dehydrogenase. The organisms were searched in PATRIC database to check their host and pathogenicity. Only the human hosts were considered and predicted to be pathogenic if involved in disease(s) according to PATRIC database [28]. We found that all the organisms are pathogenic except some species of Enterococcus such as E. hirae, mundtii, durans, saccharolyticus, villorum, gallinarum, phoeniculicola, italicus, caccae, haemoperoxidus, moraviensis, malodoratus, cecorum, raffinosus, sulfureus, avium, dispar, columbae, asini; Lactococcuspiscium and Lactococcusraffinolactis, Carnobacteriumdivergens, Staphylococcus schleiferi, Vagococcuslutrae, Melissococcusplutonius. However, literature searching helped us to conclude that Enterococcus hirae, mundtii, durans, villorum, gallinarum, italicus, cecorum, raffinosus, avium, dispar and Staphylococcus schleiferi are also involved in human diseases and in rare cases are able to transfer gene to other species [49, 50]. Rest of the bacterial species are found in animal and aquatic species [51-53]. Thus it is clearly demonstrated that *E. faecium* DO aldehyde-alcohol dehydrogenase does not share any sequence similarity with non-pathological aldehyde-alcohol dehydrogenase. Therefore, due to the above advantages, aldehyde-alcohol dehydrogenase was selected for homology modeling and subsequent structure-based drug designing.

### Homology modeling, structure validation and active site prediction

Aldehyde-alcohol dehydrogenase is a bifunctional enzymes that controls ethanol and acetate production under aerobic conditions and contains 866 residues which exhibits two separate model in SWISS-Model ranging from 18 to 453 and 459 to 866 aa representing domain acetaldehyde dehydrogenase and alcohol dehydrogenase respectively. Here, we analysed both acetaldehyde dehydrogenase and alcohol dehydrogenase domain as the template for homology modeling having sequence similarity (50.91% and 62.75% respectively) and coverages (Figure 3 & 4).

**Figure 3.**
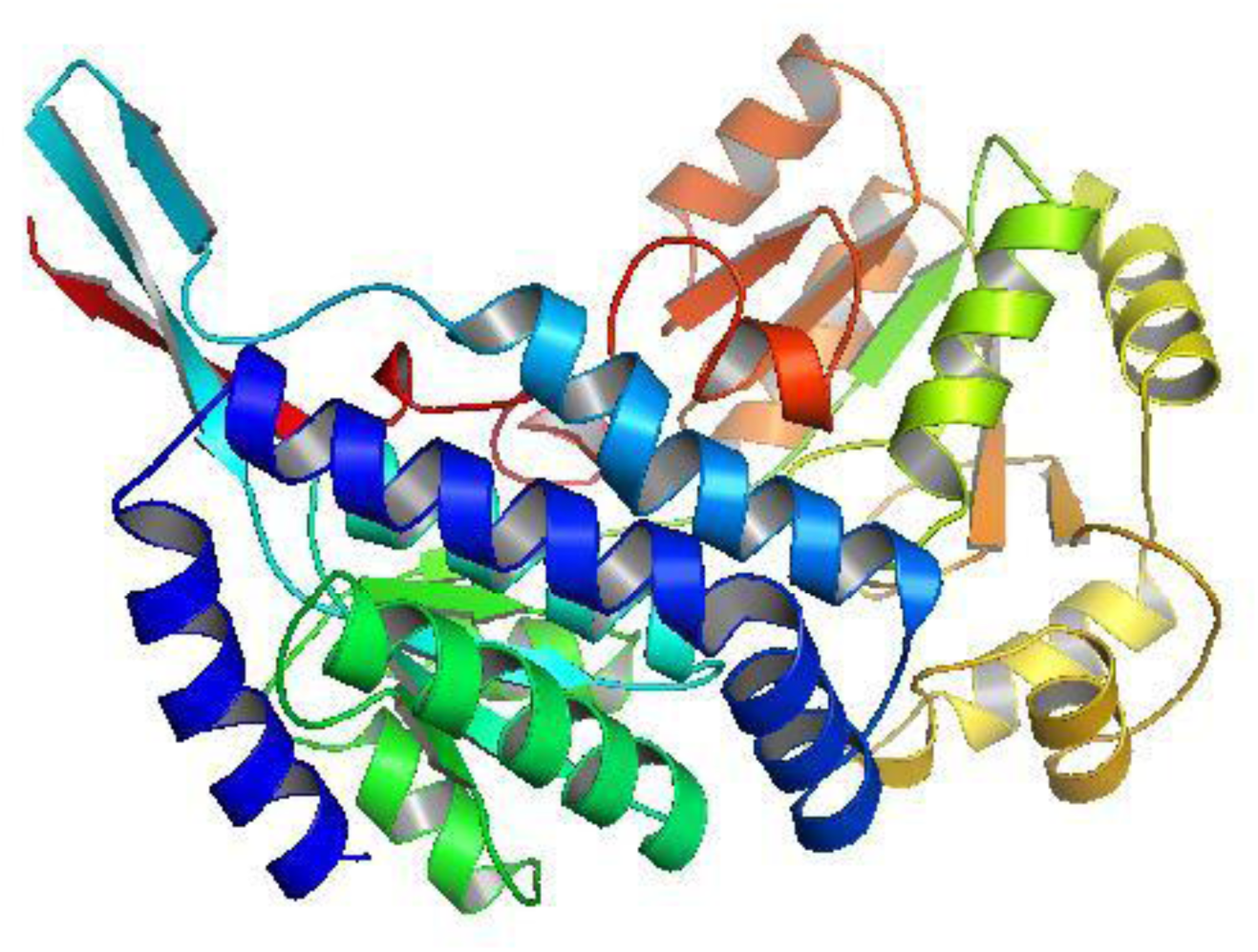
Homology modeled structure of the *E. faecium DO* acetaldehyde dehydrogenase domain by SWISS-MODEL server. **Note:** Figure generated by PyMol.

**Figure 4.**
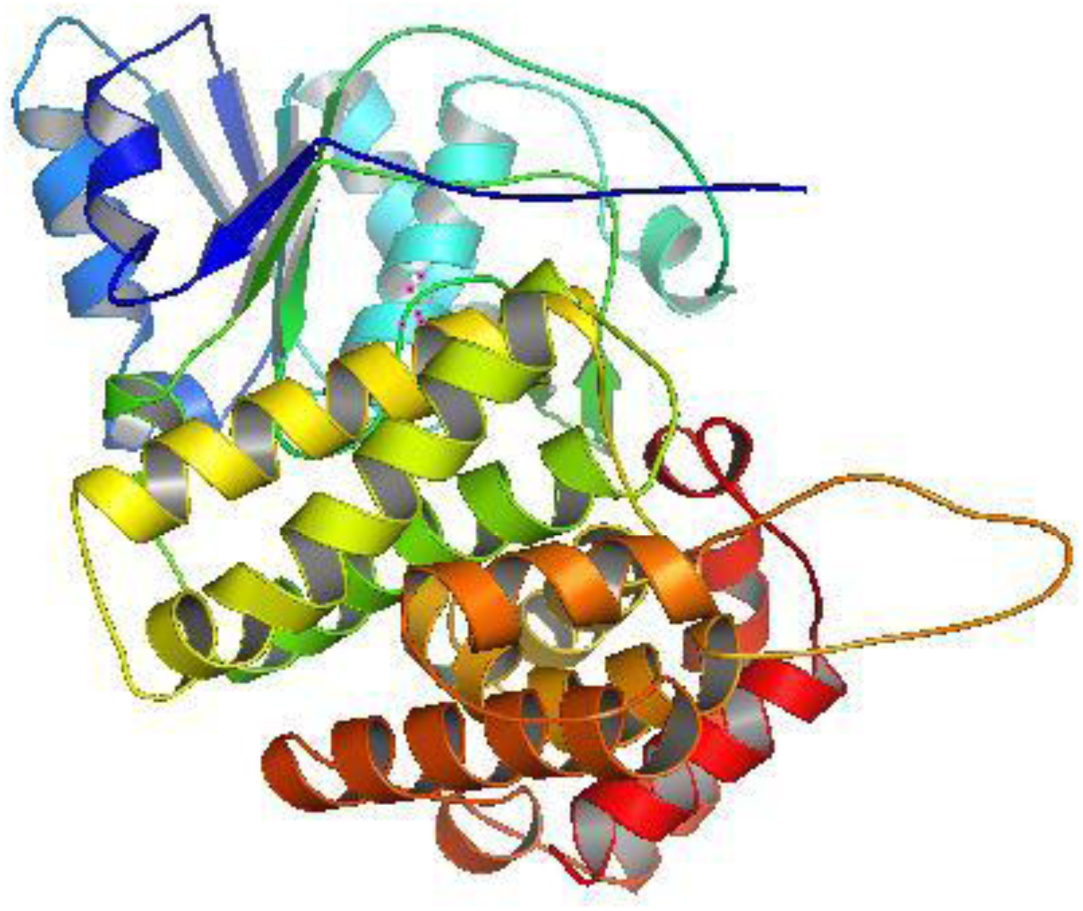
Homology modeled structure of the *E. faecium DO* alcohol dehydrogenase domain by SWISS-MODEL server. **Note:** Figure generated by PyMol.

*In silico* drug designing involving protein 3D homology modeling depends largely on the quality of the models. Psi/Phi Ramachandran plot analysis of acetaldehyde dehydrogenase built by PDBsum shows 90.5% residues resides in most favored regions, 8.2% residues in additional allowed regions and 0.8% residues in generously allowed regions (Figure 5). Similar inspection on alcohol dehydrogenase shows 90.3% residues resides in most favored regions, 8.3% residues in additional allowed regions and 0.6% residues in generously allowed regions (Figure 6). Active site is the region on the surface of a protein to which a specific substrate (ligand) or set of substrates (ligands) binds. We predicted the active sites for both acetaldehyde dehydrogenase and alcohol dehydrogenase domain by CASTp server that provides identification and measurements of surface accessible pockets for proteins with the active site residues. Largest pocket for each domain were selected for docking. The active site residues with pocket volume of 2339 and 3076.7 acetaldehyde dehydrogenase and alcohol dehydrogenase domain respectively were shown in Figure 7.

**Figure 5.**
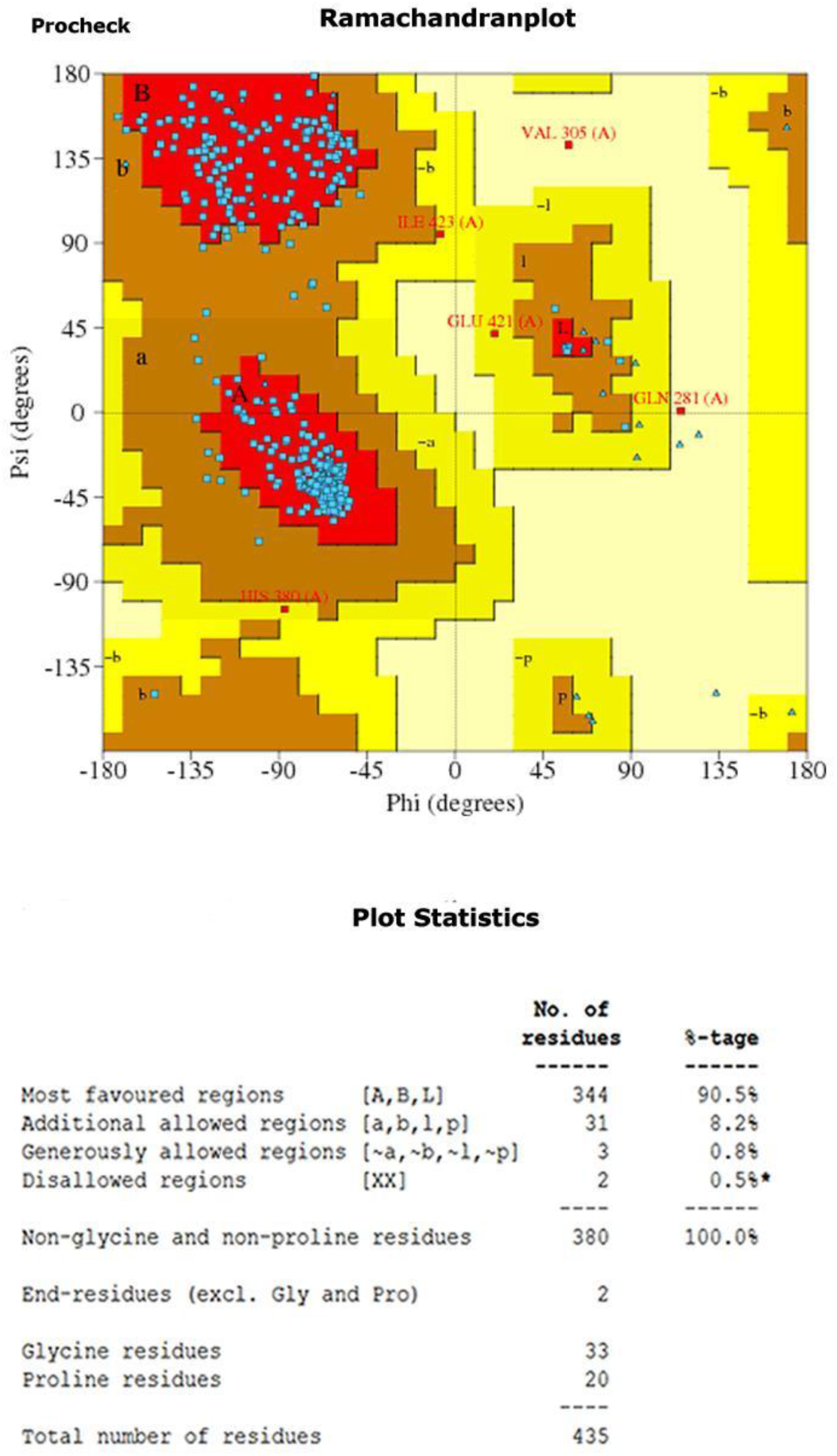
Structural validation of *E. faecium DO* acetaldehyde dehydrogenase domain. **Notes:** Result of PROCHECK verification program, showing number and percentages of residues in most favored regions (red); additional allowed regions (yellow); generously allowed regions (creamy white); and in disallowed regions (white). Based on an analysis of 118 structures of resolution of at least 2.0 angstroms and R-factor no greater than 20%, a good quality model would be expected to have over 90% in the most favored regions.

**Figure 6.**
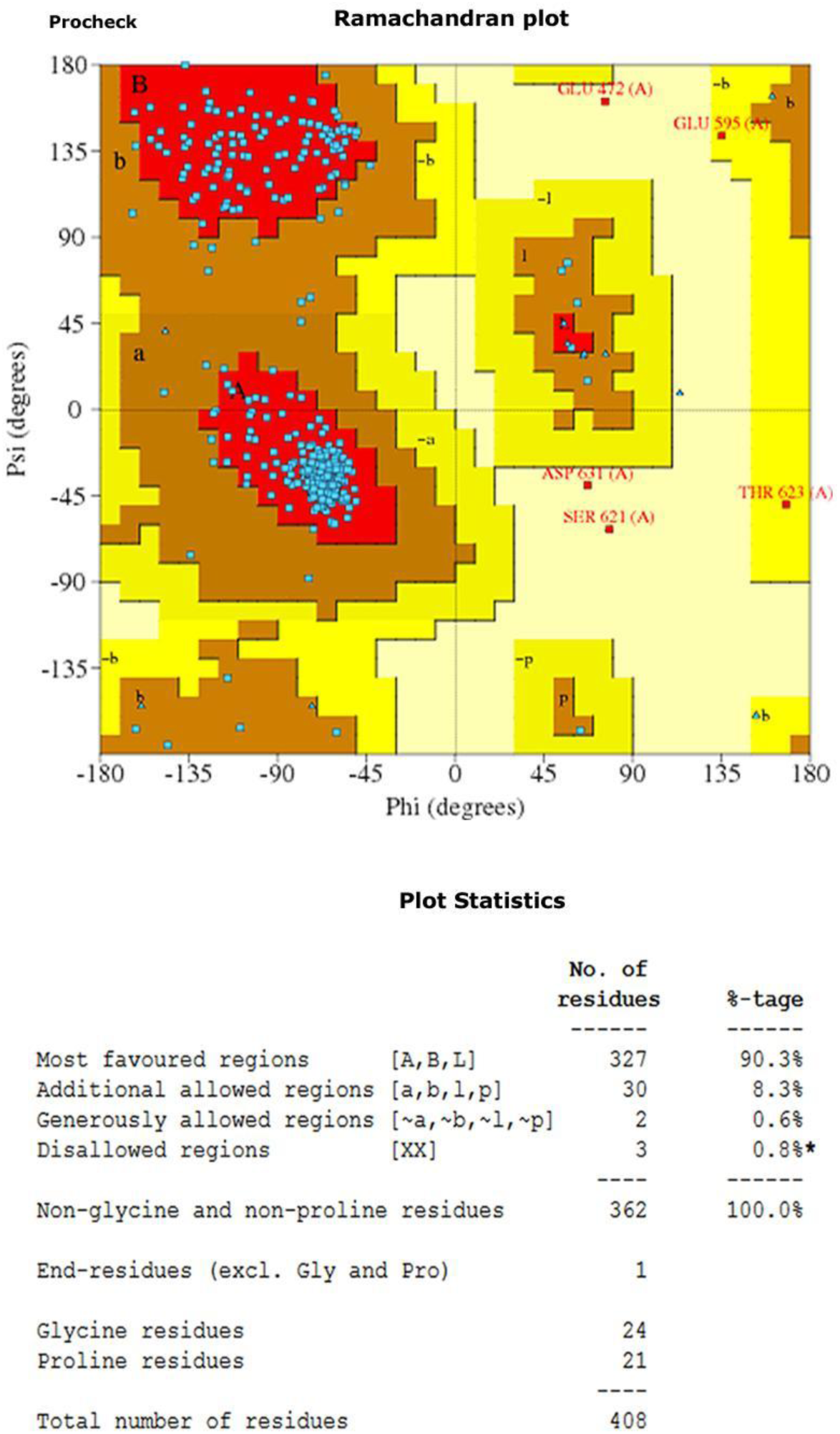
Structural validation of *E. faecium DO* alcohol dehydrogenase domain. **Notes:** Result of PROCHECK verification program, showing number and percentages of residues in most favored regions (red); additional allowed regions (yellow); generously allowed regions (creamy white); and in disallowed regions (white). Based on an analysis of 118 structures of resolution of at least 2.0 angstroms and R-factor no greater than 20%, a good quality model would be expected to have over 90% in the most favored regions.

**Figure 7.**
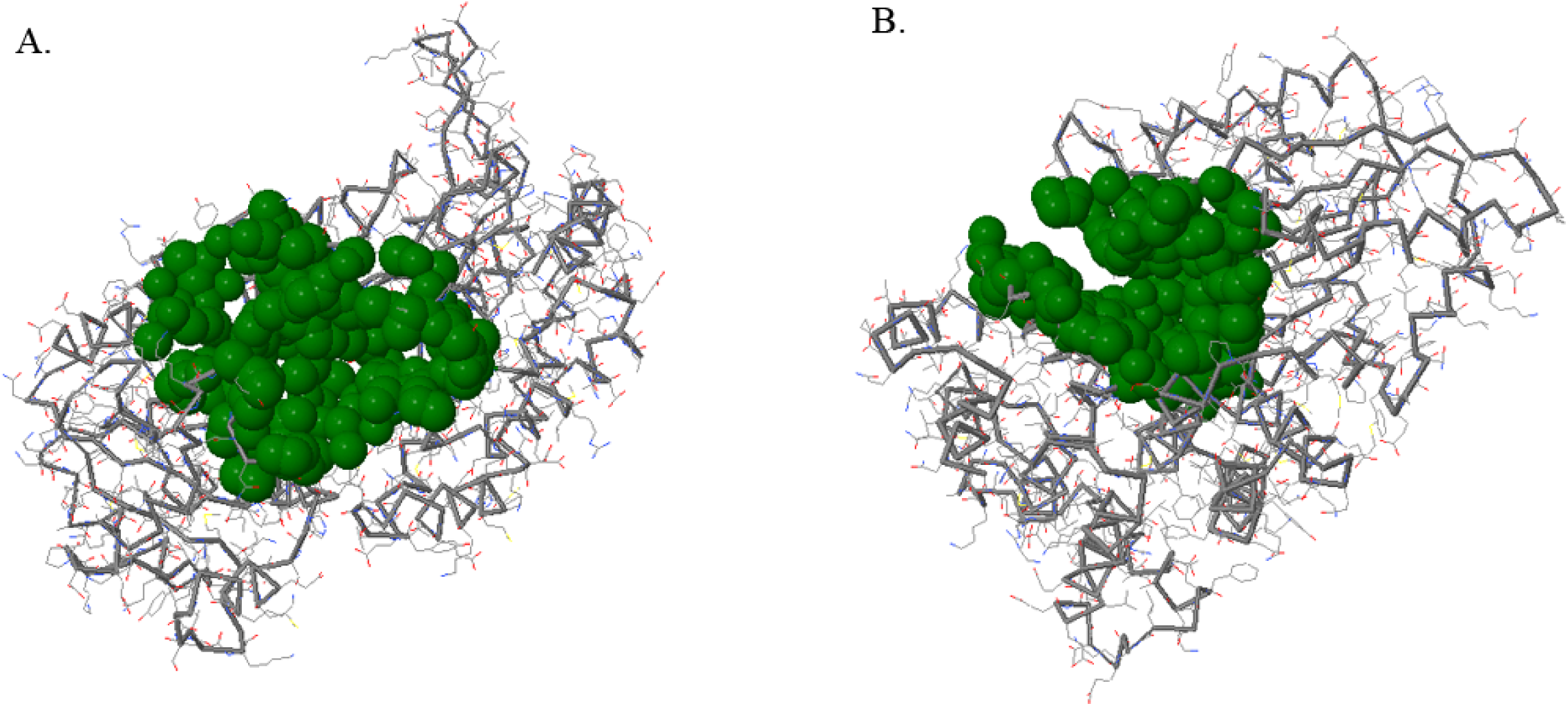
Active site prediction by CASTp. A. Active site residues (shown in green) of acetaldehyde dehydrogenase domain. B. Active site residues (shown in green) of alcohol dehydrogenase domain. **Note:** Figure prepared by CASTp server.

### Virtual screening, ADME and toxicity analysis

As Blastp search against BindingDB did not yield any significant match with our targets, we searched Protein Databank for ligand bound structure. Acetaldehyde dehydrogenase and alcohol dehydrogenase domain showed 26% (47% Positives) and 33% (49% Positives) sequence identity with Nicotinamide-adenine-dinucleotide bound PDB structure 3OX4 [54] and 4C3S [55] respectively. So we used nicotinamide-adenine-dinucleotide (NAD) as reference in virtual screening for both acetaldehyde dehydrogenase and alcohol dehydrogenase domains (Table 2). From literature search, several amino acids of active sites were found to be important for interactions with NAD in both 3OX4 and 4C3S structures (Gly98, Ser99, Thr138, Asp39, Thr139, His277, Leu187, Met42 for 3OX4; Val 221, Glu357, Ile433, Gly218, Gly236, Cys136, Cys269, Thr216, Pro135 and Thr134 for 4C3S) [54, 55]. Docking was repeated five times in order to get hits that showed stable binding energies. The top six hits showing lower binding energies were found to have stable or nearly stable binding energy and good clustering performances from the docking results (Table 3).

**Table 2:** Docking energies and important residues of the binding site observed to be interactive with the ligands, NAD.

| Domain | Compounds | Important amino acid residues involved in interactions | Docking energy (Kcal/mol) |
| --- | --- | --- | --- |
| Acetaldehyde dehydrogenase | NAD | A207, I181, T189, F149, I186, R70, P151, P225, P123, V305, V121, V310, T124, T125, G205, G206, G226, L350, E348, A304, A148, H150, M209, C257 | -8.1 |
| Alcohol dehydrogenase | NAD | V617, S610, H664, A731, K733, G735, G736, I740, T745, F615, L724, Q842, T608, D660, H742, K626, D558, H728, H732, P526, F719, N527, P628, S555, G554 | -9.1 |

**Table 3:** Lowest docking energies, important residues of the binding site observed to be interactive with the ligands, percentage of human intestinal absorption and plasma protein binding, BBB, Caco-2 cell permeability, Ames test, carcinogenicity in rats and bioavailability.

| Acetaldehyde dehydrogenase domain |  |  |  |  |  |  |  |  |  |
| --- | --- | --- | --- | --- | --- | --- | --- | --- | --- |
| Compounds | Important amino acid residues involved in interactions | Docking energy (Kcal/mol) | BBB | Caco-2 cell permeability (nm/second) | HIA % | PPB % | Carcinogenicity (Rats) | Ames test | Bioavailability |
| CID 23640940 | R70, F149, P307, P151, P225, P123, V305, V121, V310, V223, T124, T122, T125, T204, G205, G224, G206, G226, L350, E348, Y303, A304, A148, A207, H150, M209, C257, E348 | -10.8 | 0.25 | 22.64 | 88.99 | 85.97 | Negative | mutagen | Good |
| DB01369 | L350, L426, L428, T122, T125, T204, T427, C257, V223, V305, V310, V121, A304, A148, P307, P123, P151, H150, M209, G205, G206, N126, I126 | -10.3 | 0.04 | 21.27 | 93.93 | 50.76 | Negative | non-mutagen | Good |
| CID 2955941 | T125, T204, L350, L375, L426, N126, N227, V223, V305, P123, P225, P341, C257, A207, A304, G376, G206, G205, R347, Y308, E348 | -10.2 | 0.19 | 32.74 | 97.76 | 97.48 | Negative | mutagen | Good |
| DB02112 | T125, P151, F149, A148, T122, V121, M209, P307, T204, L350, T427, L428, L426, I256, N126, V223, V310, V305, P123, C257, A304, G206, G205 | -10.1 | 2.90 | 23.75 | 93.25 | 91.16 | Negative | non-mutagen | Good |
| CID 427839 | P123, P225, A207, A304, G205, G206, G208, G376, R347, Y303, V223, V305, M209, T124, T125, T204, H150, E348, L350, N126, C257 | -10.0 | 0.04 | 20.46 | 87.53 | 84.39 | Negative | non-mutagen | Good |
| CID 20910594 | S185, I186, I181, P184, P151, P225, P123, A148, T189, T204, T125, T124, F149, M209, V121, V223, H150, L350, C257, G206, G224, G205, G226, E348 | -10.0 | 0.027 | 21.07 | 96.80 | 92.8991 | Negative | non-mutagen | Good |

| Alcohol dehydrogenase domain |  |  |  |  |  |  |  |  |  |
| --- | --- | --- | --- | --- | --- | --- | --- | --- | --- |
| Compounds | Important amino acid residues involved in interactions | Docking energy (Kcal/mol) | BBB | Caco-2 cell permeability (nm/second) | Human intestinal absorption % | Plasma protein binding % | Carcinogenicity (Rats) | Ames test | Bioavailability |
| CID 16745328 | T605, N527, S610, T613, H732,G736, G739, I740, V653, A556, D493, S555, C843, K626, T657, P741, T608, H742, H728, L724, F615, F719, D558, G554 | -12.0 | 0.11 | 30.73 | 98.99 | 85.53 | positive | mutagen | Good |
| DB00872 | T605, K626, D558, S610,F615, C843, L724, F723, Q842, D656, T657, T608, D660, G743, H742, H732, F719, V617, S555, G553, G554 | -11.8 | 3.29 | 45.57 | 94.35 | 100 | positive | non-mutagen | Good |
| CID 23640943 | T605, D656, T657, P741, T608, D660, G743, I740, H742, G735, A731, H732, D493, G495, M496, P650, V649, F499, V653, K626, N527, P526, S555, G553, G554 | -11.6 | 0.10 | 20.76 | 97.66 | 100 | negative | non-mutagen | Low |
| CID 44202035 | T605, S610,T613, H732, Q842, C843,K733, K626, V625, D841, D656, T657, P741, G743, T608, V653, H742, H728, L724, H664, F615, F719, K561, V617, D558, P628, G554 | -11.4 | 2.54 | 36.80 | 96.30 | 86.50 | negative | non-mutagen | Good |
| CID 44143715 | T605, T657, P741, T608, D660, H742, H728, L724, H664, F615, F719, K626, V617, D558, N527, P528, P526, S555, G553, G554 | -11.3 | 0.03 | 18.23 | 22.90 | 73.55 | positive | non-mutagen | Good |
| CID 44144211 | F615, L724, C843, Q842, D656, T657, P741, T608, D660, H742, K626, D558, H728, H732, G743, P526, V617, F719, N527, P528, S555, G554 | -11.2 | 0.04 | 21.52 | 97.54 | 99.22 | negative | non-mutagen | Good |

Among top six molecules for acetaldehyde dehydrogenase domain, four molecules (CID 427839, CID 2955941, CID 20910594 and CID 23640940) are anti-bacterial molecules and found to be active according to PubChem Bioassay Database [56]. DB02112 (Zk-806450) is an experimental drug that belongs to the class of organic compounds known as carbazoles and Trypsin-1 is a preferred target. Interestingly, DB01369 (Quinupristin) is used in combination to treat infections by staphylococci and by vancomycin-resistant *E. faecium* that inhibits the late phase of protein synthesis [36].

For alcohol dehydrogenase domain, six molecules (CID 44143715,CID 23640943, CID 44144211, CID 44202035, CID 16745328) are active compounds in PubChem Bioassay against bacteria and fungi whereas DB00872 (Conivaptan) is an investigational drug and non-peptide inhibitor of anti-diuretic hormone. It was approved in 2004 for hyponatremia. The 2D structures and docked confirmations of top 4 molecules with receptors were shown in Figure 8. Interactions of the molecules with receptor were analysed by PoseView that included hydrogen bonds, salt bridges, metal, hydrophobic, π-π and π-cation interactions (Figure 9 and Figure 10).

**Figure 8.**
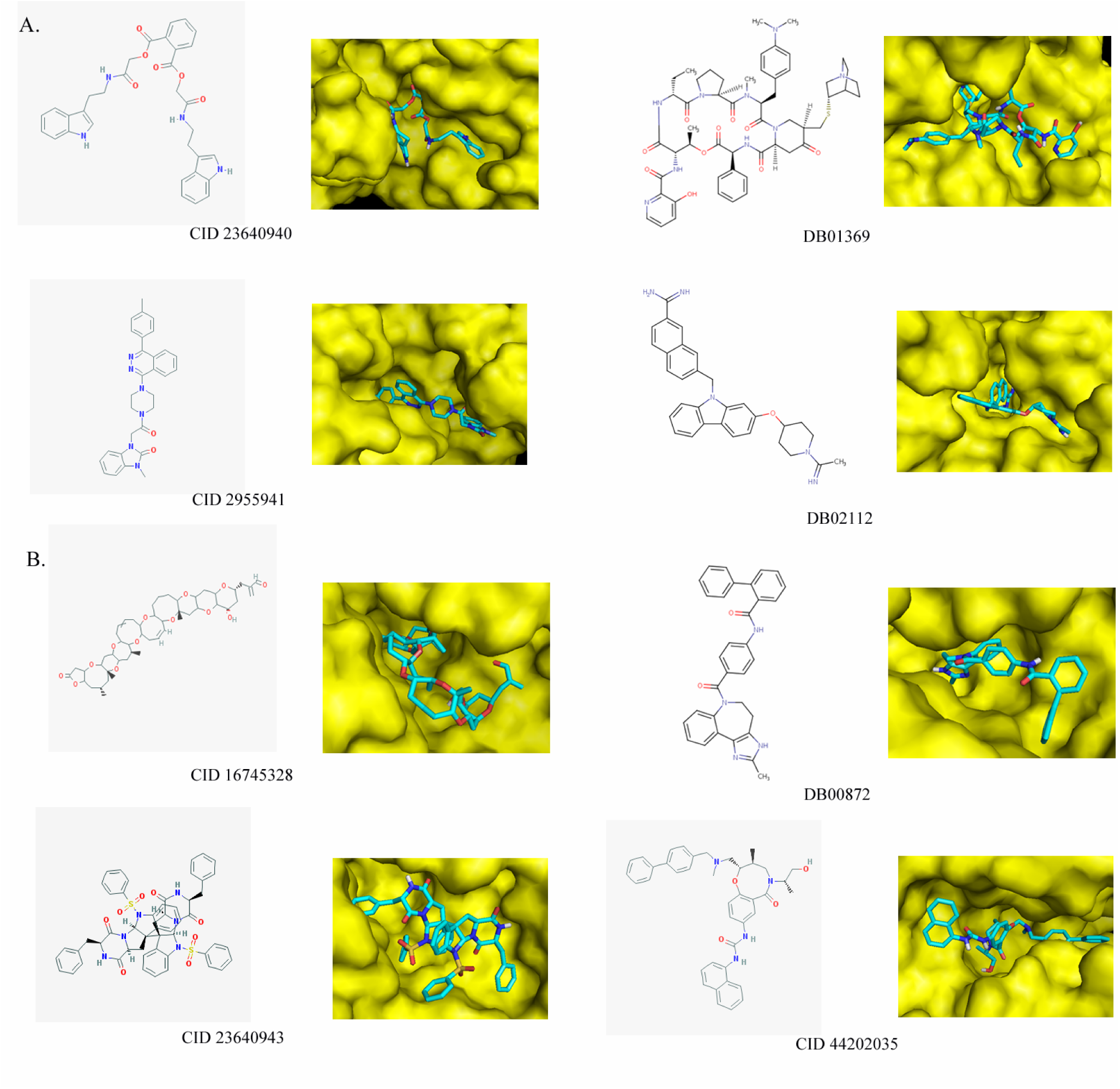
2D structures and 3D representation of docked molecules. A. Top 4 molecules of acetaldehyde dehydrogenase domain. B. Top 4 molecules of alcohol dehydrogenase domain.

**Figure 9.**
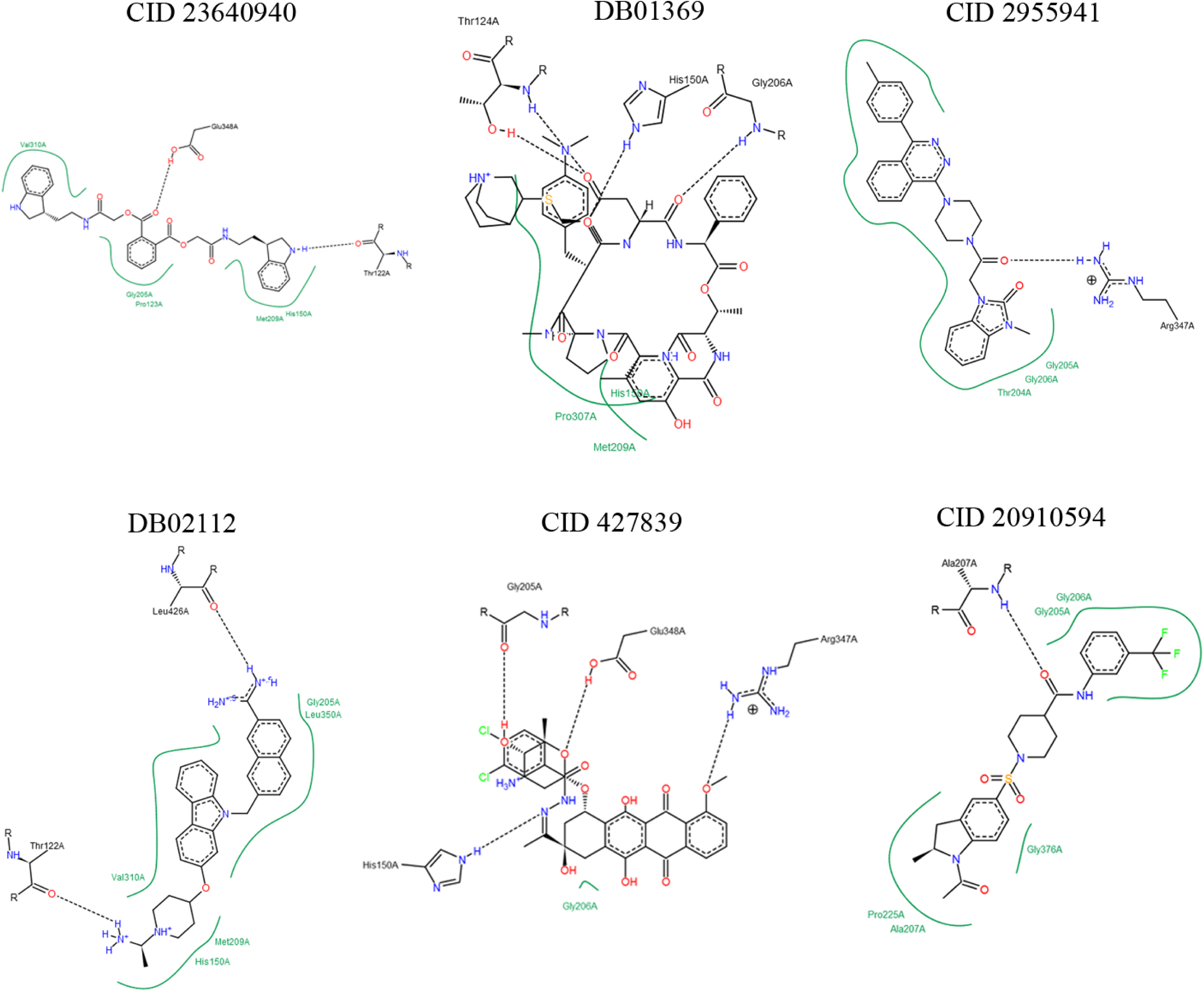
Visualization of acetaldehyde dehydrogenase domain and selected compound interactions. Note: Figure generated by PoseView software.

**Figure 10.**
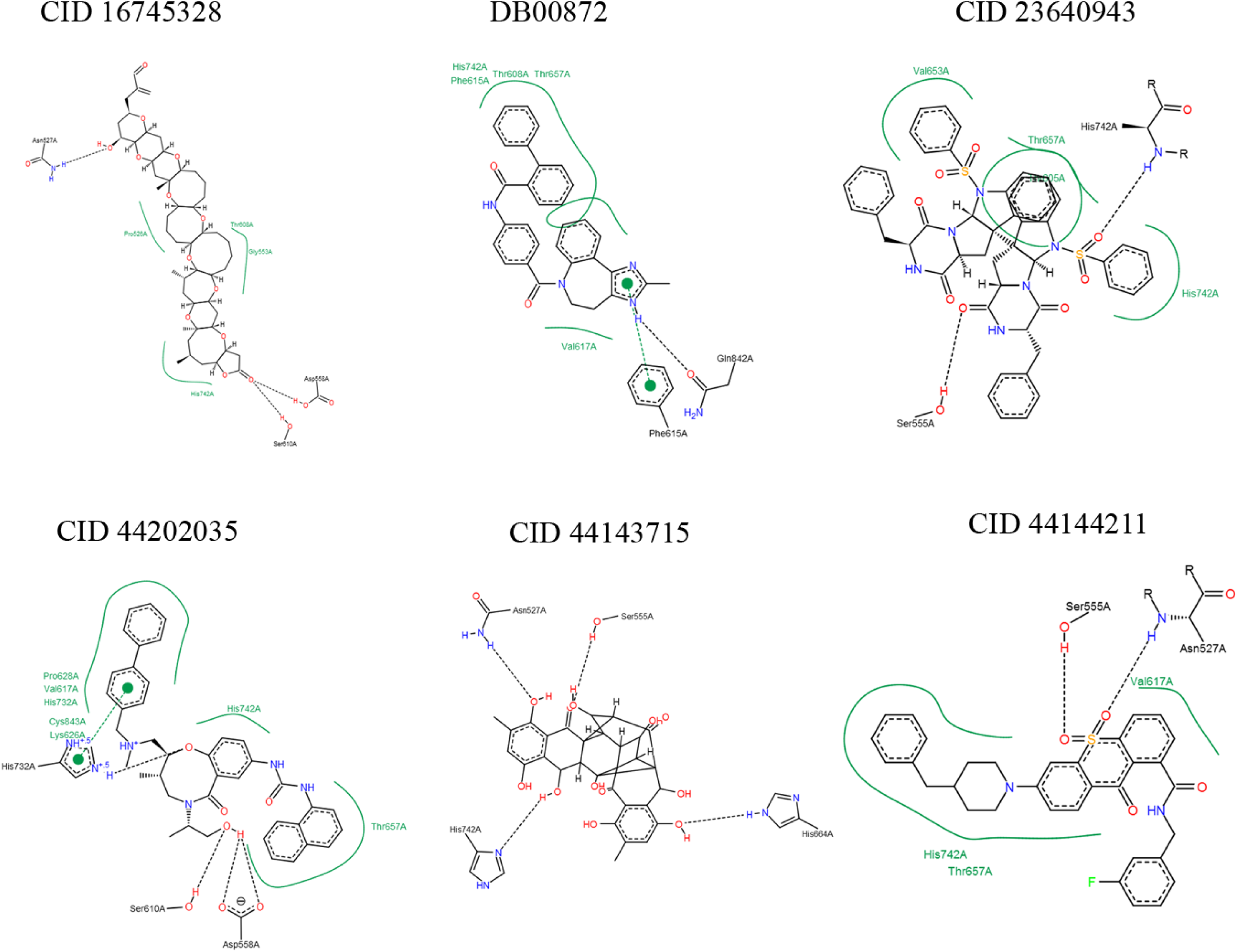
Visualization of alcohol dehydrogenase domain and selected compound interactions. Note: Figure generated by PoseView software.

ADME and Toxicity prediction were performed to identify potential candidates and undesired side effects of the molecules. For top six molecules of both acetaldehyde dehydrogenase and alcohol dehydrogenase, all molecules except CID 44143715 showed above 70% human intestinal absorption (HIA) indicating well absorbed compounds (70%∼100% according to PreADMET) that is desired for drug candidates. CID 44143715 was considered as moderately absorbed compound (20 ∼ 70 % according to PreADMET). Degree of plasma protein binding of a drug influences not only on the drug’s mode of action but also its disposition and efficacy as only the unbound drug is available for diffusion or transport across cell membranes, and for interaction with a pharmacological target. In case of acetaldehyde dehydrogenase domain, plasma protein binding prediction (%PPB) results showed that CID 23640940 and CID 427839 were weakly bound with 88.99% and 87.53% plasma protein binding whereas other molecules were strongly bound as all of them have PPB over 90% (according to PreADMET, more than 90% = Chemicals strongly bound). On the other hand, for alcohol dehydrogenase domain, three molecules each showed strong and weak binding as PPB was more than 90% and less than 90% respectively. Caco-2 cell model and MDCK cell model has been recommended as a reliable *in vitro* model for the prediction of oral drug absorption. Prediction of Caco-2 cell permeability by PreADMET fell within the range of 4-70% that was considered as middle permeability for all 12 molecules according to PreADMET.

Most of the compounds exhibited good oral bioavailability and negative result in Ames test as predicted by FAF-Drugs3 and PreADMET toxicity prediction. Ames test is a simple method to test mutagenicity of a compound. CID 23640943 showed low oral bioavailability whereas CID 23640940 and CID 16745328 were predicted as mutagen. Predicting BBB penetration means predicting whether compounds can pass across the blood-brain barrier. BBB penetration prediction by PreADMET showed that most of the compounds were found to have low and middle absorption to CNS. DB00872, CID 44202035 and DB02112 showed high absorption to CNS.

However, clear evidence of carcinogenic activity was predicted for all six compounds for acetaldehyde dehydrogenase molecules whereas no evidence of carcinogenic activity was found for 3 of alcohol dehydrogenase compounds (CID 44143715, DB00872 and CID 16745328) by PreADMET rodent carcinogenicity prediction. PreADMET predicts the result from its model which is built from the data of National Toxicology Program and US FDA. We furthermore analyzed other ADMET properties like skin permeability and P-gp inhibition, and in all of these cases results are positive for our proposed compounds (data not shown).

## Discussion

Our present study reports the first computational approach of KEGG annotated metabolic and subtractive genome analysis that led to the identification of protein that can be tested for potential drug development against *E. faecium DO*. The rationale of picking targets using computational approaches rely on searching for those genes/proteins that are absent and/or non-homologous to the host proteome but present in the pathogen. Designing a drug specific to such a target will effect only the pathogen but won’t interfere any aspects of the host biology. Therefore, using extensive *in silico* tools, we identified broad-spectrum antibiotic target and proposed aldehyde-alcohol dehydrogenase as novel drug target against highly pathogenic bacteria *E. faecium DO*.

Aldehyde-alcohol dehydrogenase is a bi-functional enzyme consisting of two domains aldehyde and alcohol dehydrogenase on its N-terminal and C-terminal respectively. This enzyme is known for ethanol and aldehyde production in anaerobic bacteria. Deletion of adhE gene encoding for this enzyme resulted in lower bacterial cell density and extended elongation period. In addition, in strains without *adhE*, changes in biochemical activity, product formation, and growth was observed which clearly indicates the association of aldehyde-alcohol dehydrogenase enzyme in regulating bacterial biological processes. Most importantly, from pathway analysis it has been found that this enzyme is involved in pathways that are responsible for antibiotic synthesis in *E. faecium DO* [57]. Since bacterial evolution aims at acquiring resistance against single or multiple antibiotics to ensure their survival in the environment, therefore, the development of new conventional antibiotics as well as novel compounds is of crying need to combat bacterial infections.

By virtual screening we identified six compounds for each domain. These molecules are analysed for their usefulness and potential side effects. Most of them showed desirable properties in all of the predictions to become potential candidates to block aldehyde-alcohol dehydrogenase of *E. faecium DO* thus hampering the survival of *E. faecium*. By *in sillico* analysis and 3D homology modeling of the two domains of aldehyde-alcohol dehydrogenase, a good quality model was generated and was verified by Ramachandhra Plot and CASTp server. A virtual screening was carried on the bioactive compounds for each domain of the protein that resulted in the identification of six and five potential novel inhibitors for aldehyde dehydrogenase and alcohol dehydrogenase domain respectively. Docking interaction analysis identified a few common active residues in the protein domains which participated in the biochemical reaction. based on their binding with each domain and ADMET properties, the shortlisted compounds are likely to interact and inhibit the replicative activity of the *E. faecium* by blocking its enzyme activity. Therefore, these ligands can be evaluated as potential anti-bacterial molecules specific to this target. However, these should be further experimentally validated for their role in bacterial survival and virulence. Such a strategy can be used to screen novel and alternative targets aiming to design new drugs and to inhibit infectious human pathogens.

## Acknowledgments

We acknowledge Taslima Haque and Sudip Biswas for cooperation and cordial support throughout this research. We would like to thank the Plant Biotechnology Lab, University of Dhaka for providing research facilities.

## References

1. Moreno, M.F., et al., The role and application of enterococci in food and health. International journal of food microbiology, 2006. 106(1): p. 1–24.

2. Holzapfel, W., et al., Enterococcus faecium SF68 as a model for efficacy and safety evaluation of pharmaceutical probiotics. Beneficial microbes, 2018. 9(3): p. 375–388.

3. Lodemann, U., et al., Effects of the probiotic Enterococcus faecium and pathogenic Escherichia coli strains in a pig and human epithelial intestinal cell model. Scientifica, 2015. 2015.

4. Bybee, S., A. Scorza, and M. Lappin, Effect of the probiotic Enterococcus faecium SF68 on presence of diarrhea in cats and dogs housed in an animal shelter. Journal of Veterinary Internal Medicine, 2011. 25(4): p. 856–860.

5. Hidron, A.I., et al., Antimicrobial-resistant pathogens associated with healthcare-associated infections: annual summary of data reported to the National Healthcare Safety Network at the Centers for Disease Control and Prevention, 2006–2007. Infection Control & Hospital Epidemiology, 2008. 29(11): p. 996–1011.

6. Control, C.f.D. and Prevention, Antibiotic resistance threats in the United States, 2013. 2013: Centres for Disease Control and Prevention, US Department of Health and ….

7. Kerschner, H., et al., Hospital outbreak caused by linezolid resistant Enterococcus faecium in Upper Austria. Antimicrobial Resistance & Infection Control, 2019. 8(1): p. 150.

8. Dubin, K. and E.G. Pamer, Enterococci and their interactions with the intestinal microbiome. Bugs as Drugs: Therapeutic Microbes for the Prevention and Treatment of Disease, 2018: p. 309–330.

9. Qin, X., et al., Complete genome sequence of Enterococcus faecium strain TX16 and comparative genomic analysis of Enterococcus faecium genomes. BMC microbiology, 2012. 12(1): p. 135.

10. Arias, C.A., et al., Cotransfer of antibiotic resistance genes and a hylEfm-containing virulence plasmid in Enterococcus faecium. Antimicrobial agents and chemotherapy, 2009. 53(10): p. 4240–4246.

11. Thurlow, L.R., et al., Enterococcus faecalis capsular polysaccharide serotypes C and D and their contributions to host innate immune evasion. Infection and immunity, 2009. 77(12): p. 5551–5557.

12. Chan, J.N., C. Nislow, and A. Emili, Recent advances and method development for drug target identification. Trends in pharmacological sciences, 2010. 31(2): p. 82–88.

13. Markwart, R., et al., The rise in vancomycin-resistant Enterococcus faecium in Germany: data from the German Antimicrobial Resistance Surveillance (ARS). Antimicrobial Resistance & Infection Control, 2019. 8(1): p. 147.

14. Ahmed, M.O. and K.E. Baptiste, Vancomycin-resistant enterococci: a review of antimicrobial resistance mechanisms and perspectives of human and animal health. Microbial Drug Resistance, 2018. 24(5): p. 590–606.

15. Mondal, S.I., et al., Identification of potential drug targets by subtractive genome analysis of Escherichia coli O157: H7: an in silico approach. Advances and applications in bioinformatics and chemistry: AABC, 2015. 8: p. 49.

16. Bhoi, P., T.K. Sahu, and A. Patel, Identification of Novel Drug Targets in Streptococcus pneumoniae using Subtractive Genomic Approach. CSVTU International Journal of Biotechnology, Bioinformatics and Biomedical, 2019. 4(3): p. 79–86.

17. Shuvo, M.S.R., S.K. Shakil, and F. Ahmed, Potential Drug Target Identification of Legionella pneumophila by Subtractive Genome Analysis: An In Silico Approach. Bangladesh Journal of Microbiology, 2018. 35(2): p. 102–107.

18. Ulitsky, I., Evolution to the rescue: using comparative genomics to understand long non-coding RNAs. Nature Reviews Genetics, 2016. 17(10): p. 601.

19. Kanehisa, M., et al., KEGG for representation and analysis of molecular networks involving diseases and drugs. Nucleic acids research, 2010. 38(suppl_1): p. D355–D360.

20. Magrane, M., UniProt Knowledgebase: a hub of integrated protein data. Database, 2011. 2011.

21. Zhang, R. and Y. Lin, DEG 5.0, a database of essential genes in both prokaryotes and eukaryotes. Nucleic acids research, 2009. 37(suppl_1): p. D455–D458.

22. Yu, N.Y., et al., PSORTb 3.0: improved protein subcellular localization prediction with refined localization subcategories and predictive capabilities for all prokaryotes. Bioinformatics, 2010. 26(13): p. 1608–1615.

23. Agüero, F., et al., Genomic-scale prioritization of drug targets: the TDR Targets database. Nature Reviews Drug Discovery, 2008. 7(11): p. 900–907.

24. Boeckmann, B., et al., The SWISS-PROT protein knowledgebase and its supplement TrEMBL in 2003. Nucleic acids research, 2003. 31(1): p. 365–370.

25. Bernstein, F.C., et al., The Protein Data Bank: a computer-based archival file for macromolecular structures. Archives of biochemistry and biophysics, 1978. 185(2): p. 584–591.

26. Hecker, N., et al., SuperTarget goes quantitative: update on drug–target interactions. Nucleic acids research, 2012. 40(D1): p. D1113–D1117.

27. Yang, H., et al., Therapeutic target database update 2016: enriched resource for bench to clinical drug target and targeted pathway information. Nucleic acids research, 2016. 44(D1): p. D1069–D1074.

28. Gillespie, J.J., et al., PATRIC: the comprehensive bacterial bioinformatics resource with a focus on human pathogenic species. Infection and immunity, 2011. 79(11): p. 4286–4298.

29. Wattam, A.R., et al., PATRIC, the bacterial bioinformatics database and analysis resource. Nucleic acids research, 2014. 42(D1): p. D581–D591.

30. Waterhouse, A., et al., SWISS-MODEL: homology modelling of protein structures and complexes. Nucleic acids research, 2018. 46(W1): p. W296–W303.

31. Arnold, K., et al., The SWISS-MODEL workspace: a web-based environment for protein structure homology modelling. Bioinformatics, 2006. 22(2): p. 195–201.

32. Melo, F. and E. Feytmans, Assessing protein structures with a non-local atomic interaction energy. Journal of molecular biology, 1998. 277(5): p. 1141–1152.

33. Laskowski, R.A., et al., PROCHECK: a program to check the stereochemical quality of protein structures. Journal of applied crystallography, 1993. 26(2): p. 283–291.

34. Guex, N. and M.C. Peitsch, SWISS-MODEL and the Swiss-Pdb Viewer: an environment for comparative protein modeling. electrophoresis, 1997. 18(15): p. 2714–2723.

35. Binkowski, T.A., S. Naghibzadeh, and J. Liang, CASTp: computed atlas of surface topography of proteins. Nucleic acids research, 2003. 31(13): p. 3352–3355.

36. Wishart, D.S., et al., DrugBank: a comprehensive resource for in silico drug discovery and exploration. Nucleic acids research, 2006. 34(suppl_1): p. D668–D672.

37. Danishuddin, M., et al., AMDD: Antimicrobial drug database. Genomics, proteomics & bioinformatics, 2012. 10(6): p. 360–363.

38. Forli, S., et al., Computational protein–ligand docking and virtual drug screening with the AutoDock suite. Nature protocols, 2016. 11(5): p. 905.

39. Lee, S., et al., The PreADME Approach: Web-based program for rapid prediction of physico-chemical, drug absorption and drug-like properties. EuroQSAR designing drugs and crop protectants: processes, problems and solutions, 2003: p. 418–20.

40. Lagorce, D., et al., FAF-Drugs2: free ADME/tox filtering tool to assist drug discovery and chemical biology projects. BMC bioinformatics, 2008. 9(1): p. 396.

41. Jimenez, A., D. Chen, and N.M. Alto, How bacteria subvert animal cell structure and function. Annual review of cell and developmental biology, 2016. 32: p. 373–397.

42. Tiwari, S., et al., Two-component signal transduction systems of pathogenic bacteria as targets for antimicrobial therapy: an overview. Frontiers in microbiology, 2017. 8: p. 1878.

43. LaRock, C.N. and V. Nizet, Cationic antimicrobial peptide resistance mechanisms of streptococcal pathogens. Biochimica et Biophysica Acta (BBA)-Biomembranes, 2015. 1848(11): p. 3047–3054.

44. Geitani, R., et al., Cationic antimicrobial peptides: alternatives and/or adjuvants to antibiotics active against methicillin-resistant Staphylococcus aureus and multidrug-resistant Pseudomonas aeruginosa. BMC microbiology, 2019. 19(1): p. 54.

45. Strong, P.J., S. Xie, and W.P. Clarke, Methane as a resource: can the methanotrophs add value? Environmental science & technology, 2015. 49(7): p. 4001–4018.

46. Petković, H., T. Lukežič, and J. Šušković, Biosynthesis of oxytetracycline by Streptomyces rimosus: past, present and future directions in the development of tetracycline antibiotics. Food technology and biotechnology, 2017. 55(1): p. 3–13.

47. Ghosal, D., et al., Current state of knowledge in microbial degradation of polycyclic aromatic hydrocarbons (PAHs): a review. Frontiers in microbiology, 2016. 7: p. 1369.

48. Barh, D., et al., In silico subtractive genomics for target identification in human bacterial pathogens. Drug Development Research, 2011. 72(2): p. 162–177.

49. Chan, T.-S., et al., Enterococcus hirae-related acute pyelonephritis and cholangitis with bacteremia: an unusual infection in humans. The Kaohsiung journal of medical sciences, 2012. 28(2): p. 111–114.

50. Wang, X., et al., High rate of New Delhi Metallo-β-Lactamase 1–producing bacterial infection in China. Clinical infectious diseases, 2013. 56(1): p. 161–162.

51. Law-Brown, J. and P.R. Meyers, Enterococcus phoeniculicola sp. nov., a novel member of the enterococci isolated from the uropygial gland of the Red-billed Woodhoopoe, Phoeniculus purpureus. International journal of systematic and evolutionary microbiology, 2003. 53(3): p. 683–685.

52. Forsgren, E., et al., Distribution of Melissococcus plutonius in honeybee colonies with and without symptoms of European foulbrood. Microbial Ecology, 2005. 50(3): p. 369–374.

53. Meslier, V., V. Loux, and P. Renault, Genome sequence of Lactococcus raffinolactis strain 4877, isolated from natural dairy starter culture. 2012, Am Soc Microbiol.

54. Moon, J.-H., et al., Structures of iron-dependent alcohol dehydrogenase 2 from zymomonas mobilis ZM4 with and without NAD+ cofactor. Journal of molecular biology, 2011. 407(3): p. 413–424.

55. Tuck, L.R., et al., Insight into Coenzyme A cofactor binding and the mechanism of acyl-transfer in an acylating aldehyde dehydrogenase from Clostridium phytofermentans. Scientific reports, 2016. 6: p. 22108.

56. Wang, Y., et al., PubChem’s BioAssay database. Nucleic acids research, 2012. 40(D1): p. D400–D412.

57. Lo, J., et al., The bifunctional alcohol and aldehyde dehydrogenase gene, adhE, is necessary for ethanol production in Clostridium thermocellum and Thermoanaerobacterium saccharolyticum. Journal of bacteriology, 2015. 197(8): p. 1386–1393.

